# Classification Of Mushrooms Using Artificial Neural Network

**DOI:** 10.1101/2022.08.31.505980

**Authors:** Aaditya Prasad Gupta

## Abstract

The artificial neural network (ANN) has had remarkable success in pattern recognition in recent years. It stands for a new learning paradigm in artificial intelligence (AI) and machine learning and has been applied to problems ranging from speech recognition to the prediction of protein secondary structure, cancers, and gene prediction. Recent breakthrough results in image analysis and speech recognition have generated massive interest in this field. However, the mathematical and computational methodology underlying deep learning models is very challenging, especially for interdisciplinary scientists. In this manuscript, a Neural Network model is used to classify whether a given mushroom is edible or poisonous using Tensorflow in Python based on the attributes present in the dataset. The dataset includes descriptions of hypothetical samples corresponding to 23 species of gilled mushrooms in the Agaricus and Lepiota Family Mushroom, drawn from The Audubon Society Field Guide to North American Mushrooms (1981).

## 1 Introduction

A mushroom is the fleshy, spore-bearing fruiting body of a fungus, typically produced above ground, on soil, or on its food source which is both edible and poisonous. The edible mushrooms include nutritional content and health benefits. However, some mushroom species are toxic and contain poisonous substances that could cause illness and lead to death. Mushroom poisoning accounts for approximately 70% of natural poisoning and often causes death^1^. However, there are only 30-50 poisonous species among the thousands of species found on earth, and of these, no more than 10 are fatally poisonous. Eating mushrooms collected in the wild is risky and should only be undertaken by individuals knowledgeable in mushroom recognition^2^. The mutual best practice for the wild mushroom pickers is to focus on gathering a small number of visually distinct, edible mushroom species that cannot be simply confused with poisonous varieties^3^. Distinguishing edible from poisonous mushroom species needs meticulous care to detail; there is no single trait by which all toxic mushrooms can be recognized, nor one by which all edible mushrooms can be recognized. The figure 1 shows the cartoon picture of the different types of mushrooms.

**Figure 1.**
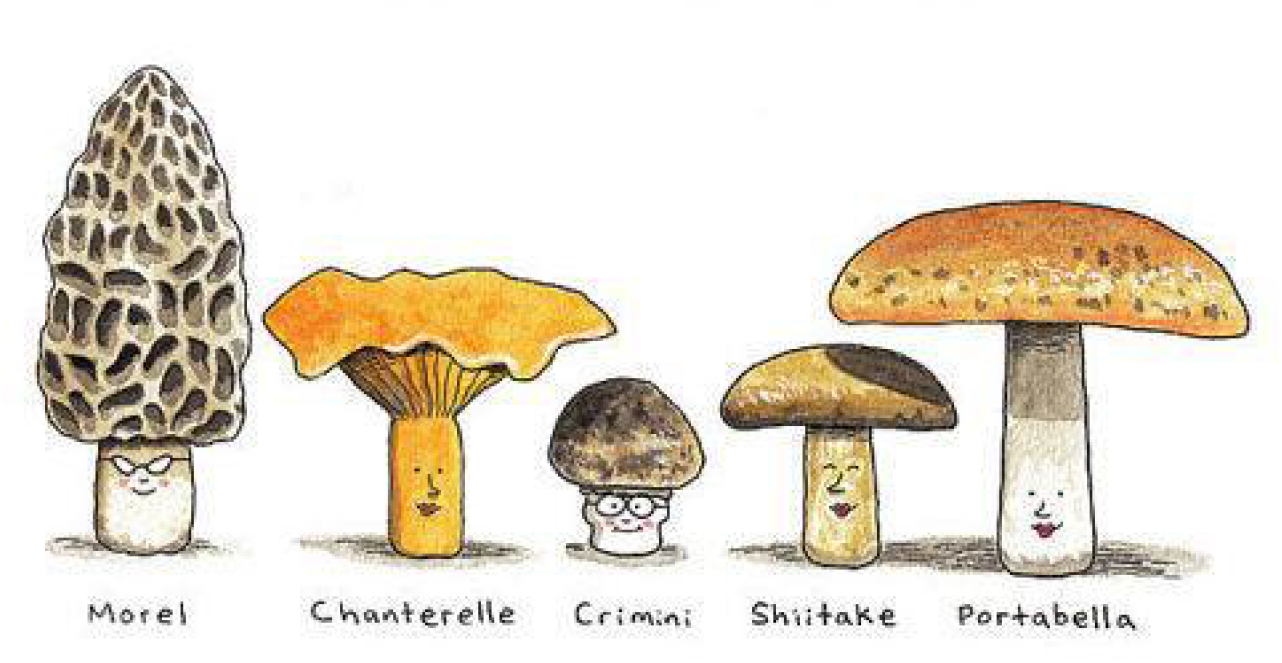
Different types of mushroom. The image is retrieved from https://www.kaggle.com.

The Artificial Neural Network system (ANNs) can be able to classify if a mushroom is poisonous or not, using data mining as one of the approaches for obtaining computer-assisted knowledge. Here, we apply the Neural Network model to the mushroom hunting dataset that contains 8124 instances, 22 attributes, and 2 possible classes of mushrooms to classify it as. edible or poisonous. Our approach is to classify or predict the class of mushrooms using a forward-feed supervised Neural Network using the different attributes based on Tensorflow^4^ in Python. This manuscript is arranged in the following way to incorporate the information about both mushrooms and Artificial Neural Networks, and is inspired by the article^5,6^. In section 2 we discuss about the short description of Artificial Neural Network. In subsection 2.1 the application of ANNs in the field of emerging biology is discussed. Similarly, in section 3 we discuss about the classification of mushrooms using the ANNs whereas in subsection 3.2 and 3.1 we discuss about the method used for classification and detailed information about the data. While section 4 discusses about the results. Finally, this manuscript ends with a discussion and a brief conclusion in section 5.

## 2 Artificial Neural Network

Artificial Neural Networks (ANNs), usually simply called neural networks (NNs) or neural nets, are computing systems inspired by the biological neural networks that constitute animal brains. Warren McCulloch and Walter Pitts^7^ (1943) opened the subject by creating a computational model for neural networks^8^. Later, In 1958, psychologist Frank Rosenblatt invented the perceptron, the first artificial neural network^9–11^, funded by the United States Office of Naval Research^12^. ANN is configured for solving artificial intelligence problems without creating a model or real biological system^13^. They are based on computers and man’s brain abilities. Similarly, the main asset of the neural network is the ability of their neurons to take part in analysis while working simultaneously but independently from each other. Artificial neural networks are also good at analyzing large sets of unlabeled, often high-dimensional data-where it may be difficult to determine a prior which questions are most relevant and rewarding to ask.

Artificial Neural Networks are a sub-set of machine learning and are the heart of deep learning algorithms. A standard neural network (NN) consists of many simple, connected processors called a neuron, each producing a sequence of real-valued activation. An artificial neuron receives a signal and then processes them and can signal neurons connected to it. ANNs are comprised of node layers, containing an input layer, one or more hidden layers, and an output layer. Each node or artificial neuron, that connects to another has an associated weight and threshold. If the output of any individual node is above the specified threshold value, that node is activated, sending data to the next layer of the next layer of the network. The equation representing feed-forward multi-layer ANN is shown in equation 1. The figure 2 shows a pictorial diagram of an Artificial Neural Network, with an input layer, a hidden layer, and an output layer. The input layer has n inputs, while the hidden layer has n nodes, whereas the Network has n distinct outputs. All the neurons are densely connected. Figure 3 shows the chart flow for the neural network along with the weights, activation functions, and the relations between the inputs to each neuron.

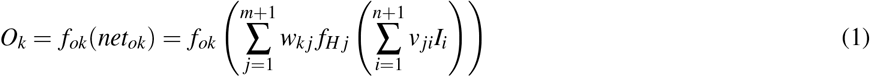

**Figure 2.**
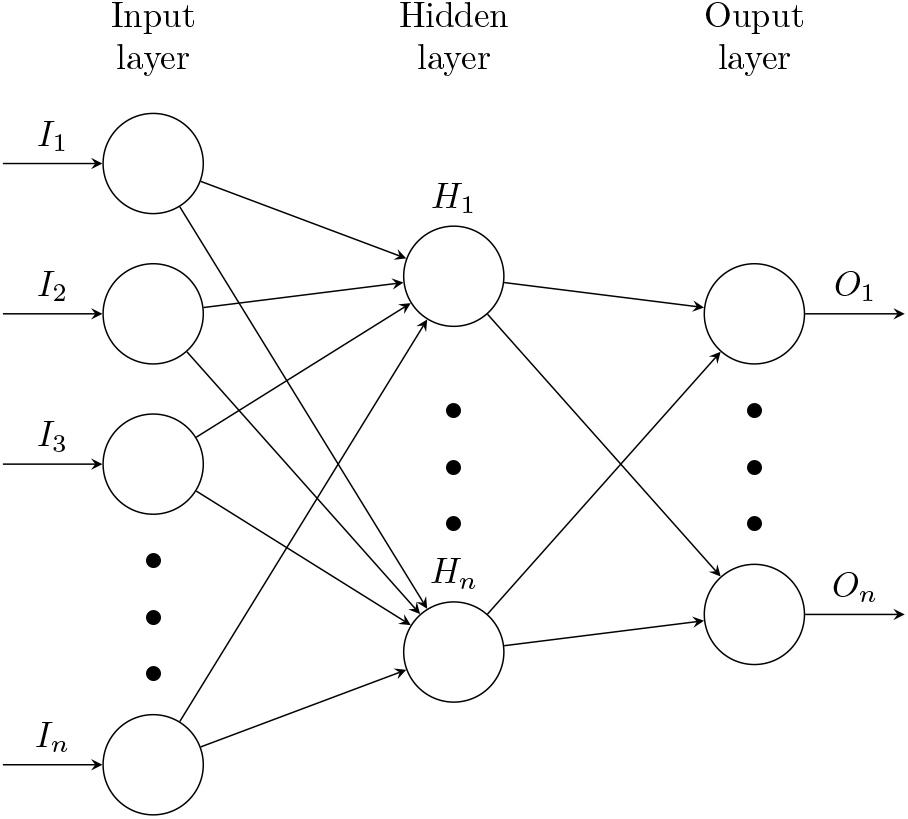
Artificial Neural Network with different layers. The figure is drawn in Latex^14^ using tikz package^15^.

**Figure 3.**
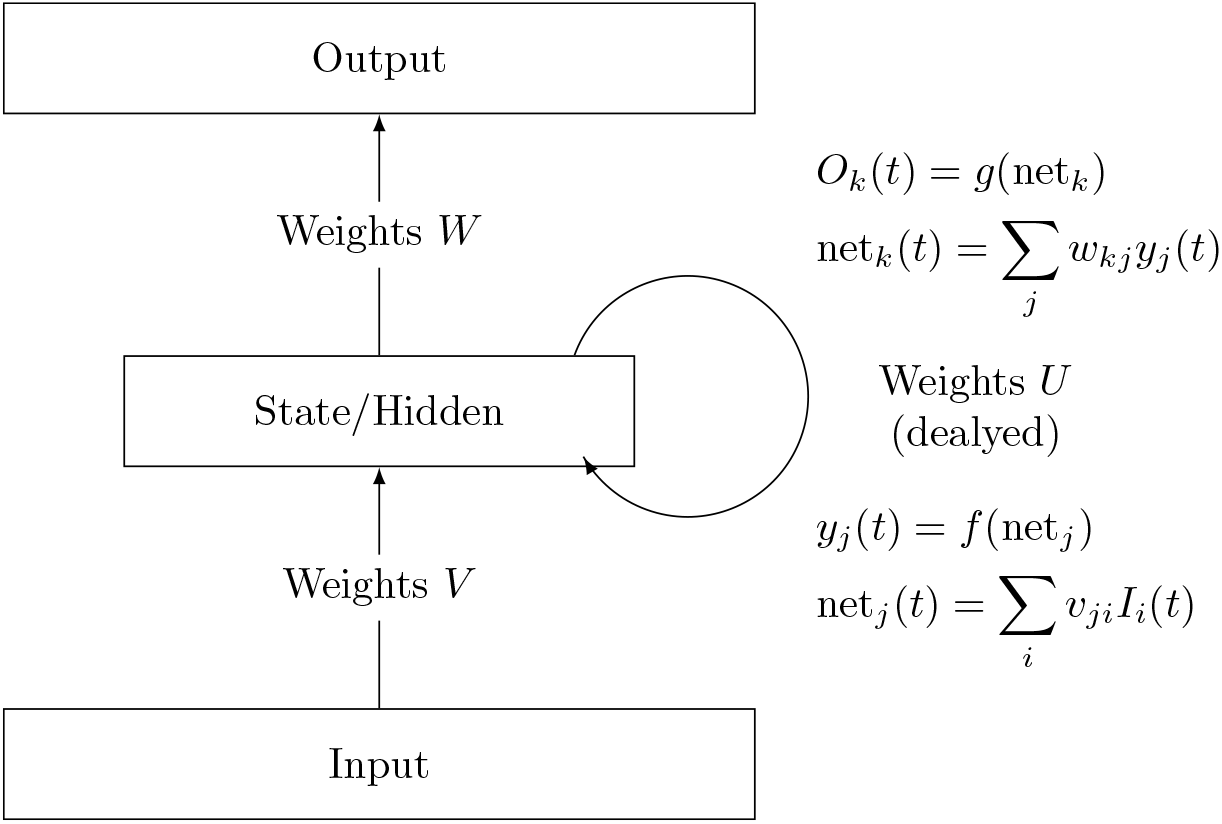
Artificial Neural Networks flowchart. The figure is drawn in Latex using tikz package.

Here, *I*_*i*_ is the *i*^*th*^ input on the input layer, *v* _*ji*_ is the respective weight. The *w*_*k* *j*_ is the weight on the node from the hidden layer to the output layer, *f*_*H* *j*_ is the corresponding activation function, *O*_*k*_ is the target output, and *f*_*ok*_ is the respective activation function.

### 2.1 Application in Biology

The neural network has been applied widely in biology since the 1980s^16^, and has advanced tremendously in the field of research, and others lately^17^. Baldi and Brunak used application in biology to explain the theory of neural network^18^. The spectrum of applications of an artificial neural network is very wide. ANNs is used for the diagnosis of different diseases in both plant and animal caused due to different factors^19^. The capacity of ANNs to analyze large amounts of data and detect patterns warrants application in the analysis of medical images, classification of tumors, and prediction of survival^20^ have somehow made easy research in medical biology. It is also very useful for solving the problem of any disease which has many confusing symptoms^21^.

Neural networks have also been actively used in many bioinformatics applications such as DNA sequence, prediction, protein secondary structure prediction, gene expression profiles classification, and analysis of gene expression patterns^22–24^. The concepts of neural network used in pattern classification and signal processing gene processing have been successfully applied in bioinformatics^25^. Neural Likewise, in medical science ANNs have been extensively applied in diagnosis, electronic signal analysis, radiology, etc. ANNs have been used by many authors for clinical research. The application of ANNs is increasing in medical data mining^26^. In agriculture, Artificial neural networks are one of the most popular tools for high production efficiency combined with high-quality products^27^. ANNs can replace the classical methods of modeling many issues, and are one of the main alternatives to classical mathematical models^28^. Neural Networks have also been applied to the analysis of gene expression patterns as an alternative to hierarchical cluster methods. Neural networks have been used in agriculture for the selection of appropriate net for a plant during sudden and quick changes in environmental conditions and predict the result of it^29^.

## 3 Classification of Mushroom

Mushrooms are fungi that are cherished for their flavor as well as for their nutritional value. They are low in salt and sugar and are a rich natural source of vitamin D.Some mushrooms are edible while some are poisonous which may cause an illness. So classification is required. It has different parts such as cap, scale, gills, tubes, pores, ring, stipe, stalk, scales, and volva as shown in figure 4. The cap is the top of the mushroom (and often looks sort of like a small umbrella). Mushroom caps can come in a variety of colors but most often are brown, white, or yellow. Gills, Pores, or Teeth appear under the mushroom’s cap. They look similar to a fish’s gills. The ring (sometimes called the annulus) is the remaining structure of the partial veil after the gills have pushed through. The stem or stipe is the tall structure that holds the cap high above the ground. The volva is the protective veil that remains after the mushroom sprouts up from the ground. As the fungus grows, it breaks through the volva. Spores are microscopic seeds acting as reproductive agents; they are usually released into the air and fall on a substrate to produce a new mushroom.

**Figure 4.**
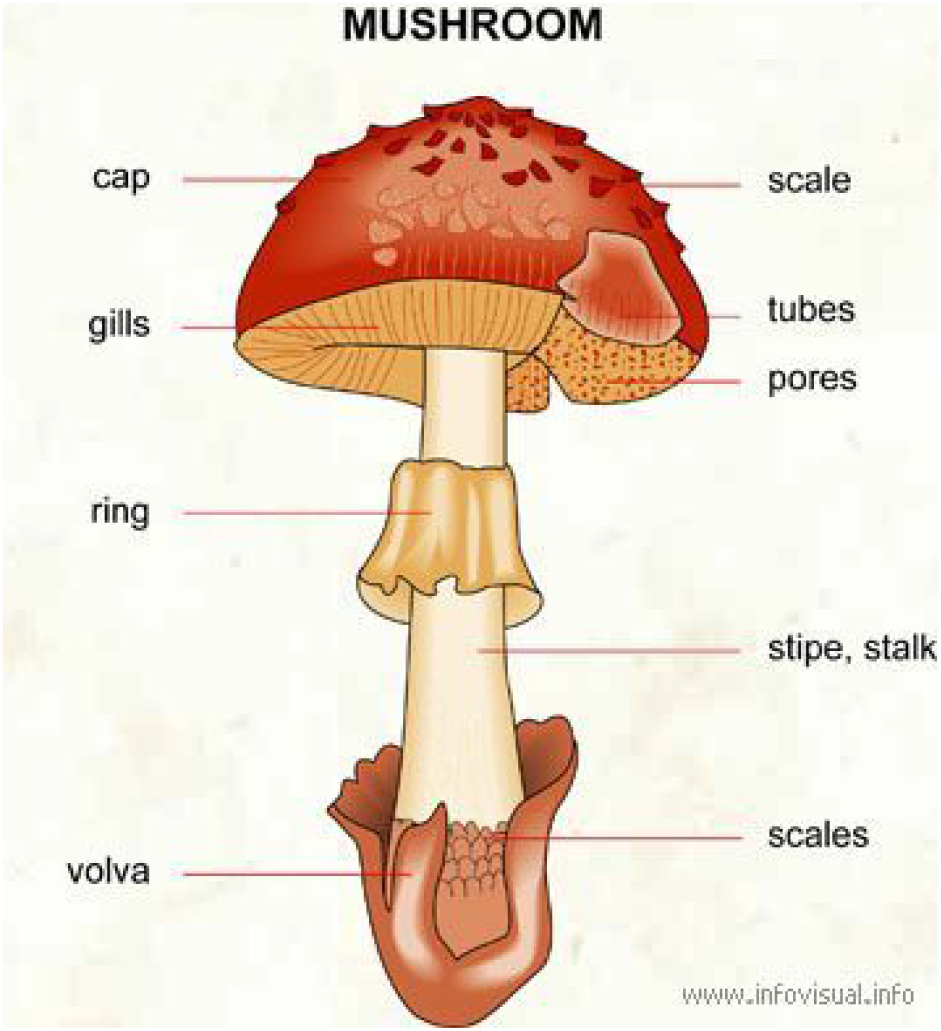
Labelled diagram of a mushroom. The image is retrieved from https://www.kaggle.com.

### 3.1 Data

This dataset was originally contributed to the UCI Machine Learning repository nearly 30 years ago, mushroom hunting (otherwise known as “shrooming”) and is retrieved from UCI Machine learning repository^30^. This dataset consists of 8124 instances, 22 attributes, and 2 possible classes. It helps to learn which features (inputs) spell certain death and which are most palatable in this dataset of mushroom characteristics. This dataset includes descriptions of hypothetical samples corresponding to 23 species of gilled mushrooms in the Agaricus and Lepiota Family Mushroom drawn from The Audubon Society Field Guide to North American Mushrooms (1981). Each species is identified as definitely edible, definitely poisonous, or of unknown edibility and not recommended. This latter class was combined with the poisonous one. The Guide clearly states that there is no simple rule for determining the edibility of a mushroom; no rule like “leaflets three, let it be” for Poisonous Oak and Ivy.

The 22 attributes that the dataset contains are:

1. **cap-shape:** bell=b, conical=c, convex=x, flat=f, knobbed=k, sunken=s.
2. **cap-surface:** fibrous=f, grooves=g,scaly=y, smooth=s.
3. **cap-color:** brown=n, buff=b, cinnamon=c, gray=g, green=r, pink=p, purple=u, red=e, white=w, yellow=y.
4. **bruises:** bruises=t, no=f.
5. **odor:** almond=a, anise=l, creosote=c, fishy=y, foul=f, musty=m, none=n, pungent=p, spicy=s.
6. **gill-attachment:** attached=a, descending=d, free=f, notched=n.
7. **gill-spacing:** close=c, crowded=w, distant=d.
8. **gill-size:** broad=b, narrow=n.
9. **gill-color:** black=k, brown=n, buff=b, chocolate=h, gray=g, green=r, orange=o, pink=p, purple=u, red=e, white=w, yellow=y.
10. **stalk-shape:** enlarging=e, tapering=t.
11. **stalk-root:** bulbous=b, club=c, cup=u, equal=e, rhizomorphs=z, rooted=r, missing=?.
12. **stalk-surface-above-ring:**fibrous=f, scaly=y, silky=k, smooth=s.
13. **stalk-surface-below-ring:** fibrous=f, scaly=y, silky=k, smooth=s.
14. **stalk-color-above-ring:** brown=n, buff=b, cinnamon=c, gray=g, orange=o, pink=p, red=e, white=w, yellow=y.
15. **stalk-color-below-ring:** brown=n, buff=b, cinnamon=c, gray=g, orange=o, pink=p, red=e, white=w, yellow=y.
16. **veil-type:** partial=p, universal=u.
17. **veil-color:** brown=n, orange=o, white=w, yellow=y.
18. **ring-number:** none=n, one=o, two=t.
19. **ring-type:** cobwebby=c, evanescent=e, flaring=f, large=l, none=n, pendant=p, sheathing=s, zone=z.
20. **spore-print-color:** black=k, brown=n, buff=b, chocolate=h, green=r, orange=o, purple=u, white=w, yellow=y.
21. **population:** abundant=a, clustered=c, numerous=n, scattered=s, several=v, solitary=y.
22. **habitat:** grasses=g, leaves=l, meadows=m, paths=p, urban=u, waste=w, woods=d.

In figure 5, blue line denotes poisonous where as buff color denotes edible mushrooms. Most of the mushrooms which have pendant or flaring ring types are edible of if they have large ring type, it is most likely to be poisonous. Similarly, in figure 6, the pie charts show the spore print color percentages in dataset according to class of mushrooms. If we see the edible mushrooms, 41.4% of edible mushrooms have brown spore print and 39.2% have black spore print color. Likewise, if we see the pie chart of poisonous mushrooms we find that 40.4% of poisonous mushrooms have chocolate print and 45.3% have white spore print color. In figure 7, the mushrooms having red and orange stalk color counts above ring are likely to be edible while buff and brown stalk color are poisonous. The mushrooms which have red, orange and gray stalk color counts below ring are mostly edible where as buff and brown are poisonous.

**Figure 5.**
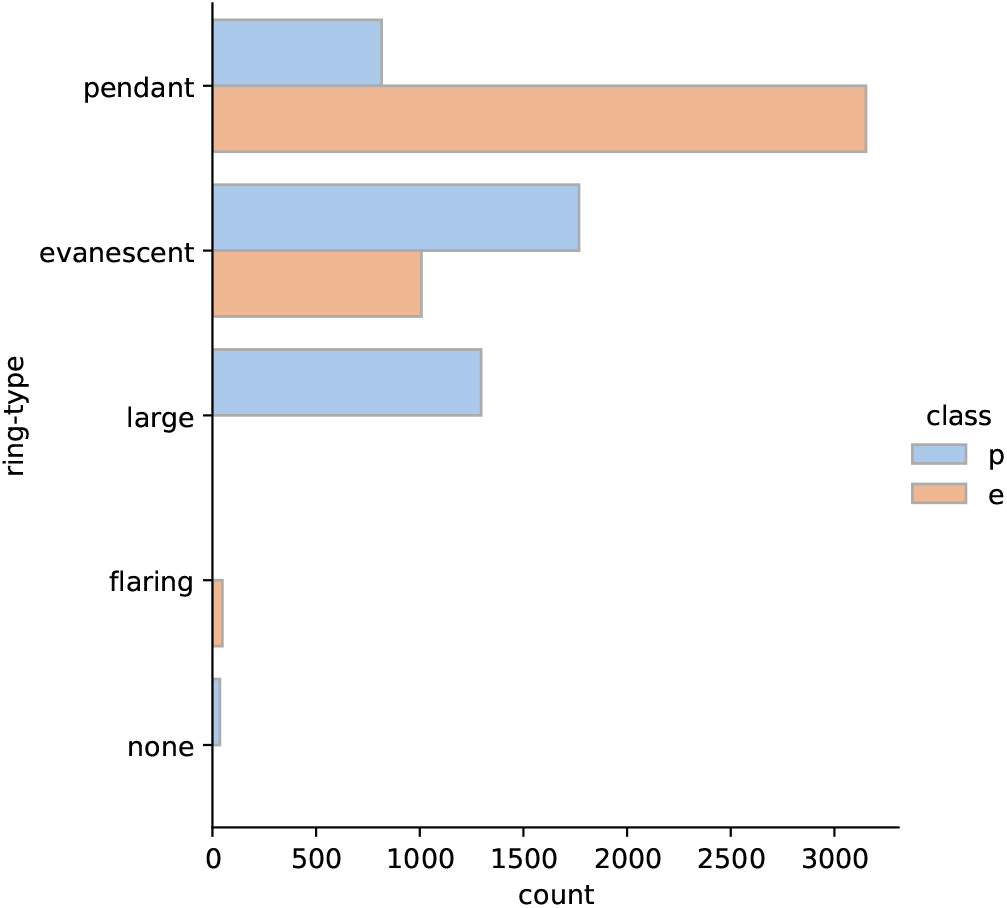
Ring type

**Figure 6.**
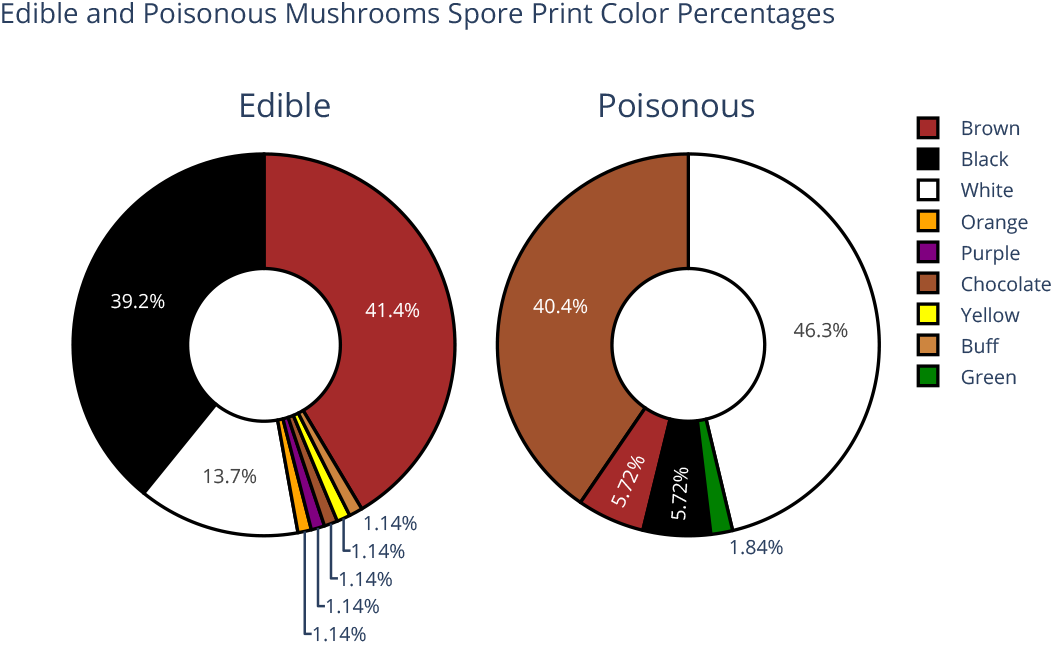
Spore print

**Figure 7.**
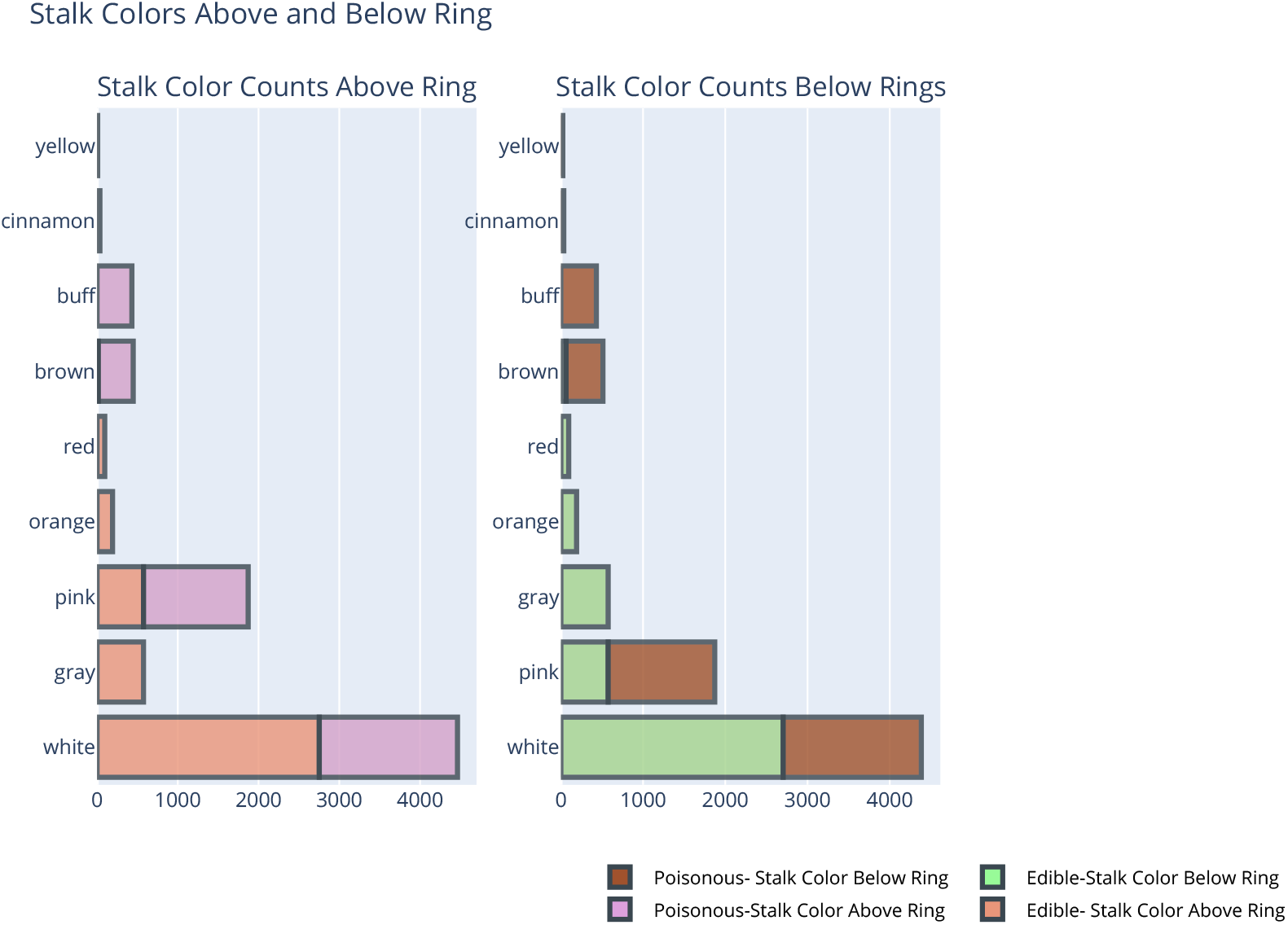
Stalk Color

### 3.2 Methodology

In this, we have separated the input and target and scaled all the inputs. We created a Neural Network model for the classification of mushrooms using Tensorflow in Python. The dataset was split into training and testing purposes to control the over-fitting. We minimized the loss and changed the hyper-parameters of the model to acquire the best fit results of the model as well as prevent it from over fit. The detail of the process involved in the Neural Network training is shown in figure 8.

**Figure 8.**
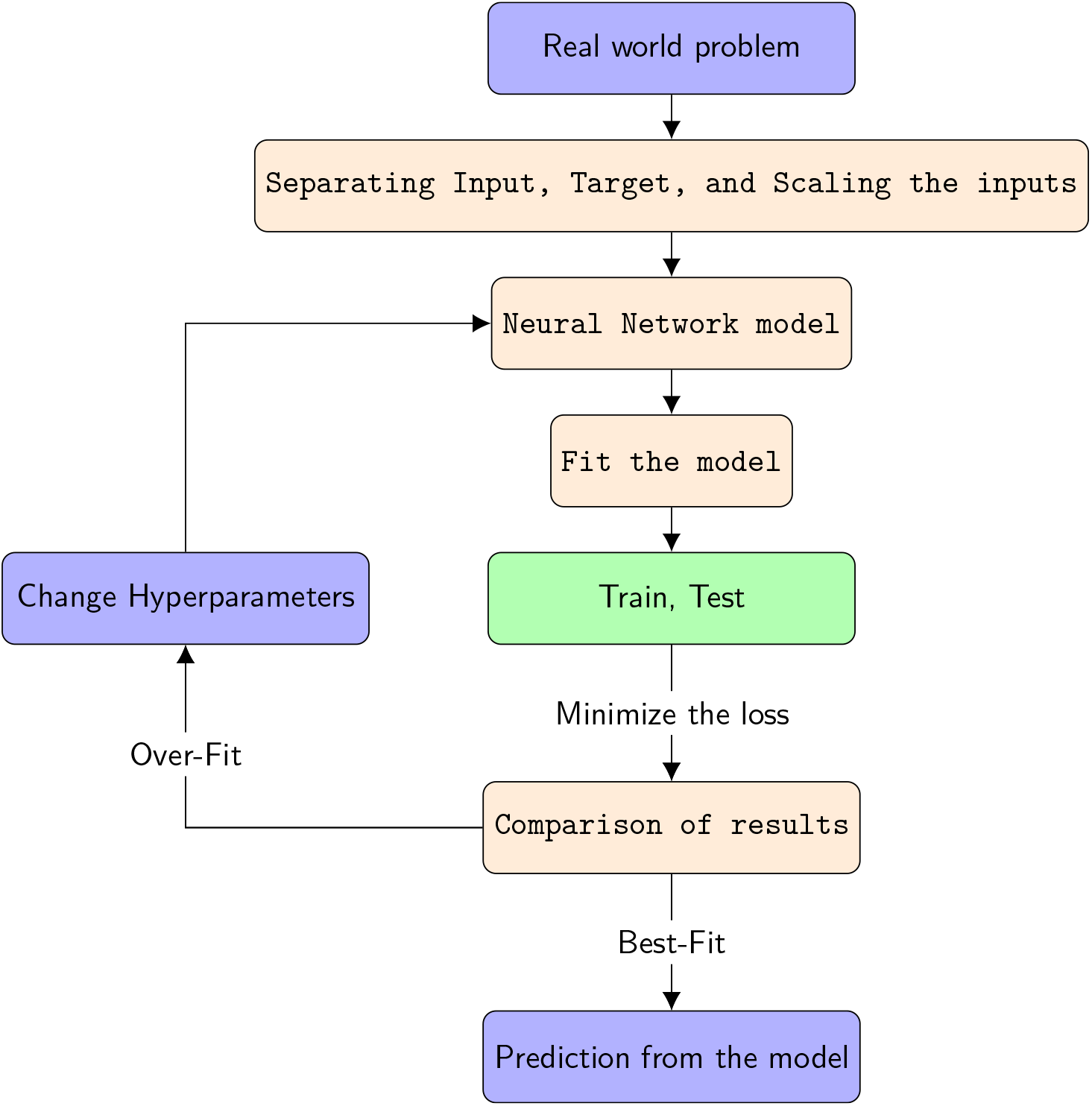
Steps involved in Mushroom classification using Artificial Neural Network. The figure is drawn in Latex using tikz package.

The learning process for an ANN is the process through which the weights of the network are determined. This is achieved by adjusting the weights until certain criteria are satisfied. The weights of the ANN are adjusted iteratively such that the difference between the actual output of the ANN and the target is minimized. The most common supervised learning method is based on the gradient descent learning rule. The method optimizes the network weights such that a certain objective function E is minimized by calculating the gradient of E in the weight space and moving the weight vector along the negative gradient. The binary cross-entropy (BCE) is used for training^31^. The BCE calculates the loss by computing the following average as shown in equation 2.

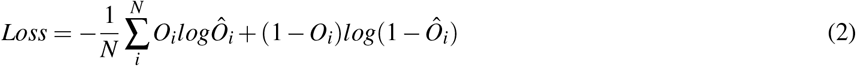

where *Ô*_*i*_ is the i-th scalar value in the output, *O*_*i*_ is the corresponding target value, and N is the number of scalar values in the model output.

For each iteration (usually called epoch), the gradient descent weight optimization contains two phases:

1. feed-forward pass in which the output of the network is calculated with the current value of the weights, activation function, and bias.
2. backward propagation in which the errors of the output signal are propagated back from the output layer towards the input layer and the weights are adjusted as a function of the back-propagated errors.

## 4 Results

A Multi-layer Neural Network acts as a classifier to distinguish signal from background for the classification of mushroom and is constructed in python using Tensorflow. The weights of input are also generated using a random number generator in python. A Neural Network model with 16 nodes, 1 discriminator layer with adam optimizer, and binary cross-esntropy was used for optimization and loss calculation. The small learning avoids overshooting.

Here, figure 9 shows the neural network output distribution for the training and testing sample of mushrooms. In separation, we have a target 1 and 0. The output that tends to 1 is poisonous near 1, while the one that tends to 0 is edible. Here the Neural Network seems to classify the mushroom as poisonous or edible with a large degree of separation. To measure the extent of separation, further curves such as ROC, loss, and accuracy are plotted.

**Figure 9.**
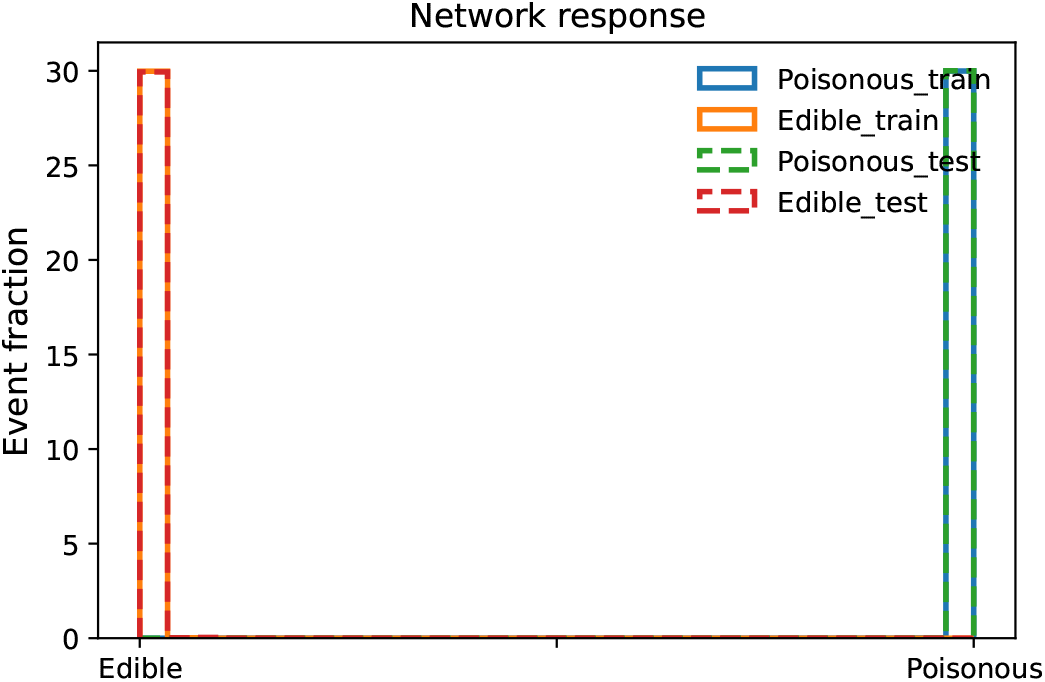
Output of the ANNs for the the train and test sample

The figure 10 shows the ROC curve of the neural network output. ROC curve, also known as the Reciever Operating Characteristics curve, is a metric used to measure the performance of a classifier model. The ROC curve depicts the rate of true positives with respect to the rate of false positives, therefore highlighting the sensitivity of the classifier model. Here, from the area under the ROC curve, it is clear that the efficiency of our neural network training is 1 which is perfect and there is no overfitting of data. The above figure 11 shows the loss of the neural network setup with varying the number of epochs. Loss is a number indicating how bad was a single example. We need a loss function to measure how good our prediction model does in terms of being able to predict the expected outcome (or value). If the model prediction is perfect, the loss is zero. Here, both the training and testing sample decreases at once with the increase in the number of epochs which shows that there is no loss or zero loss and the model prediction is perfect. In figure 12, it shows the accuracy of the neural network setup with varying the number of epochs. Accuracy is the number of true positives and true negatives divided by the number of true positives, true negatives, false positives, and false negatives. The accuracy tells overall how often the model is making a correct prediction. Here, both the training and testing sample increases at the same path with an increase in the number of epochs which shows that the accuracy of the model is 100%.

**Figure 10.**
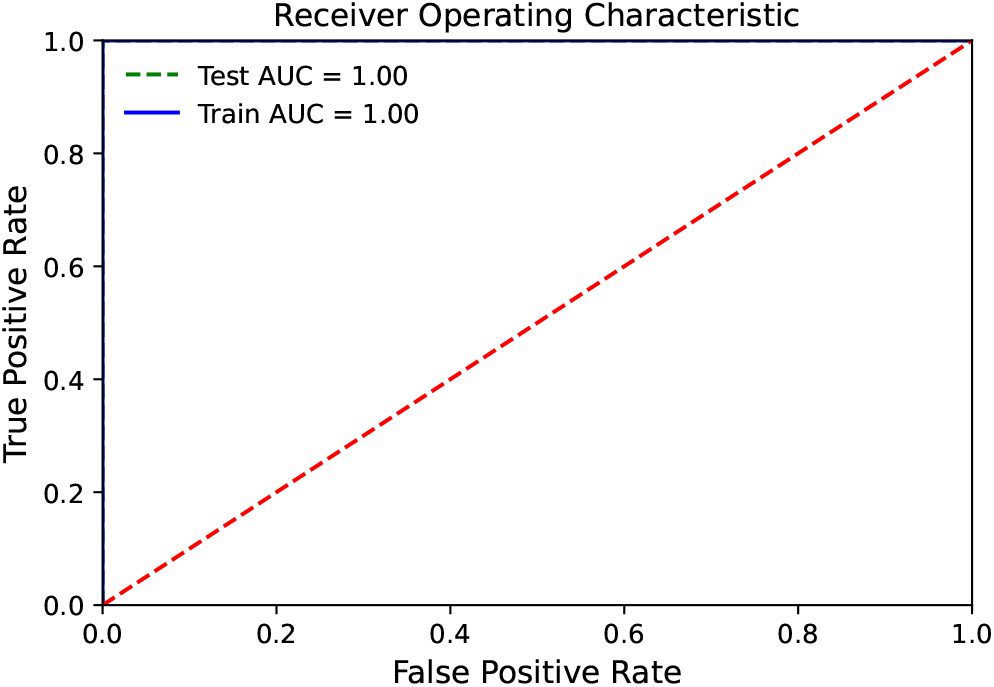
ROC curve for the train and test data sample

**Figure 11.**
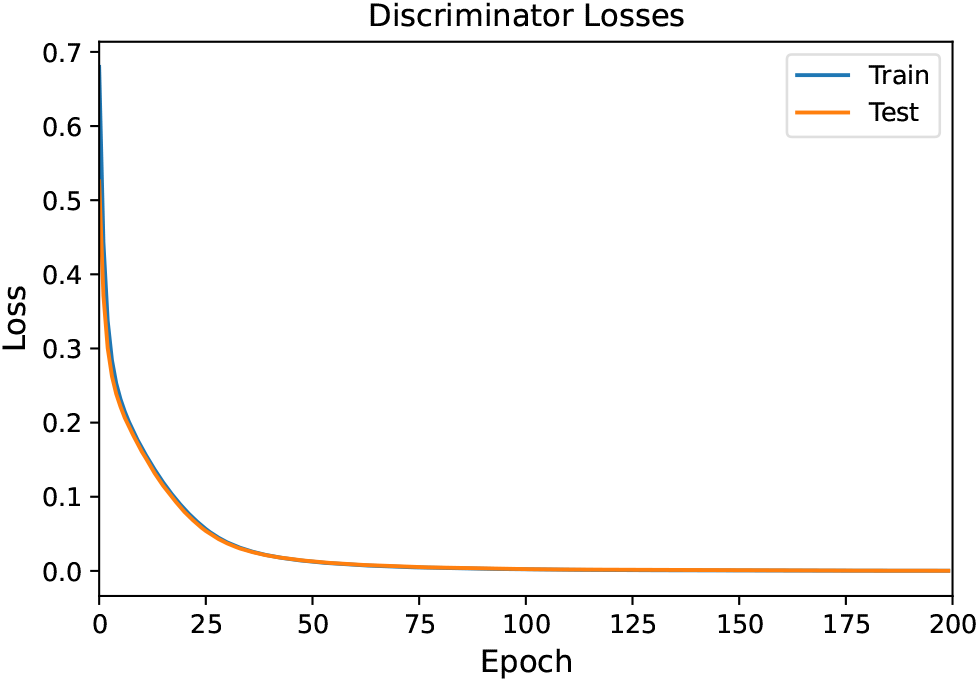
Losses of the Neural Network configuration with number of iterations for train and test sample

**Figure 12.**
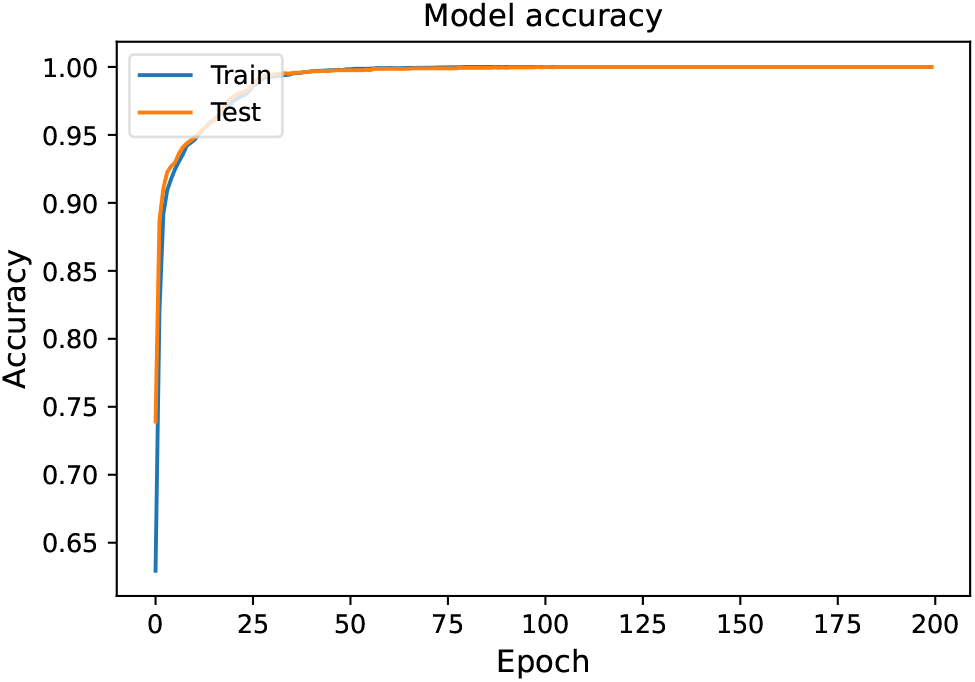
Accuracy of the Neural Network model for the train and test samples

## 5 Conclusion and Discussion

A Multi-layer Neural Network as a classifier to distinguish the mushroom as edible or poisonous was constructed. Python modules were used which have many excellent and well-maintained libraries that facilitate high-level scientific computing analyses. The modules such as numpy^32^ offer a numerical processing library that supports multi-dimensional arrays, sklearn^33^ which offers various features such as classification, regression, and clustering algorithms including support-vector machines, random forests gradient boosting, etc, matplotlib^34^ which offers a plotting library with tools to display data in a variety of ways, plotly^35^ which offers interactive online graphing, analytics, and statics tools, as well as scientific graphic libraries used in anaconda software^36^.

A Python-based feed-forward multi-layer perceptron with one input layer, one hidden layer, and one output layer (Neural Network) using Tensorflow was trained for this purpose using hypothetical samples that correspond to 23 species of gilled mushrooms in the Agaricus and Lepiota Family Mushroom, drawn from The Audubon Society Field Guide to North American Mushrooms (1981. We used the prediction power of a neural network to classify whether a mushroom is edible or poisonous. It was found that for the higher number of epochs and smaller rate, the networks perform better. Our network achieved an accuracy of over 99%. The average predictability rate was above 99% for the prediction of whether the mushroom is edible or poisonous. This study showed that Artificial Neural Network is capable of predicting a large amount of input given for the classification of mushroom whether it is edible or poisonous.

